# *Dehalobium* species implicated in 2,3,7,8-tetrachloro-*p*-dioxin dechlorination in the contaminated sediments of Sydney Harbour Estuary

**DOI:** 10.1101/2021.12.12.472303

**Authors:** Matthew Lee, Gan Liang, Sophie I. Holland, Casey O’Farrell, Keith Osbourne, Michael J. Manefield

**Affiliations:** UNSW Water Research Centre, School of Civil and Environmental Engineering, UNSW, Sydney, NSW 2052, Australia; Coffey Tetra Tech, Southbank Victoria 3006, Australia; Environment Protection Science Branch, Department of Planning, Industry & Environment, Lidcombe, NSW, 2141

## Abstract

Polychlorinated dibenzo-*p*-dioxins and furans (PCDD/F) are some of the most environmentally recalcitrant and toxic compounds. They are naturally occurring and by-products of anthropogenic activity. Sydney Harbour Estuary (Sydney, Australia), is heavily contaminated with PCDD/F. Analysis of sediment cores revealed that the contamination source in Homebush Bay continues to have one of the highest levels of PCDD/F contamination in the world (5207 pg WHO-TEQ g^-1^) with >50% of the toxicity attributed to 2,3,7,8-tetrachlorodibenzo-*p*-dioxin (2,3,7,8-TCDD) the most toxic and concerning of the PCDD/F congeners. Comparison of congener profiles at the contamination source with surrounding bays and historical data provided evidence for the attenuation of 2,3,7,8-TCDD and other congeners at the source. This finding was supported by the detection of di-, mono- and unchlorinated dibenzo-*p*-dioxin. Microbial community analysis of sediments by 16S amplicon sequencing revealed an abundance of lineages from the class Dehalococcoidia (up to 15% of the community), including the genus *Dehalobium* (up to 0.5%). Anaerobic seawater enrichment cultures using perchloroethene as a more amenable growth substrate enriched only the *Dehalobium* population by more than six-fold. The enrichment culture then proved capable of reductively dechlorinating 2,3,7,8-TCDD to 2,3,7-TCDD and octachlorodibenzo-*p*-dibenzodioxin to hepta and hexa congeners. This work is the first to show microbial reductive dehalogenation of 2,3,7,8-TCDD with a bacterium from outside the *Dehalococcoides* genus, and one of only a few that demonstrates PCDD/F degradation in a marine environment.

## Introduction

Poly chlorinated dibenzo-*p*-dioxins and furans (PCDD/F) are ubiquitous and persistent organic pollutants. Their ubiquity is due to their production by a number of both natural and anthropogenic processes. The most significant process being combustion of organic matter such as in wildfires and volcanic eruptions^1^. Anthropogenic combustion events such as waste and fuel (wood and coal) burning also contribute to PCDD/F global background concentrations^2, 3^. PCDD/F have long been viewed as threat to human and environmental health because they are highly toxic, lipophilic and resistant to metabolism, resulting in bioaccumulation and biomagnification^4^. 2,3,7,8-tetrachlorodibenzo-*p*-dioxin (2,3,7,8-TCDD) is one of the most toxic substances known with an LD^50^ of 0.6 μg.kg^-1^ in guinea pigs^5, 6^. The PCDD/F toxicity mode is through binding to the aryl hydrocarbon receptor (AHR). The degree of toxicity of each congener correlates with its binding affinity for the AHR^7^. Through this phenomenon toxicological equivalency factors (TEFs) were assigned to the 17 most problematic PCDD/F congeners (i.e. seven PCDDs and 10 PCDF)^8^. TEFs range in orders of magnitude difference, the highest being 2,3,7,8-TCDD (TEF = 1) and the lowest being octachlorodibenzo-*p*-dioxin (OCDD, TEF = 0.0003)^8^. The toxicity equivalent (TEQ) is the toxicity-weighted mass of each individual congener. In 1998 the World Health Organization (WHO) adopted the TEF approach as a means to quantify PCDDF/F and compare toxicity in environments containing congener mixtures^4^.

Sydney Harbour Estuary includes the waters of the Sydney Harbour, Middle Harbour and the Parramatta River, and is the centerpiece of the city of Sydney, the most populated city in Australia (>5 million people). The estuary has been the site of heavy industrial activity since the early 1800s^9^. One notable industry that has left an indelible mark on the estuary was a chemical manufacturing site on Homebush Bay, approximately 15 km west of the estuaries entrance. Homebush Bay is a 0.8 km^2^ embayment opening into the Parramatta River. For more than 50 years Union Carbide operated a facility that synthesized pentachlorophenol (PCP, a timber preservative) and the herbicides 2,4-dichloro and 2,4,5-trichlorophenoxyacetic acid (2,4-D and 2,4,5-T)^10^. By-products from these syntheses include: Octa, hepta and hexa congeners of PCDD/F from PCP, and 2,3,7,8-TCDD from 2,4-D and 2,4,5-T^5, 11^. Prior to 1970 PCDD/F waste from the facility was landfilled nearby and contaminated soil was then used to reclaim land from the eastern side of bay, eventuating in massive contamination of the bays sediments^10^.

The most recent PCDD/F concentration data for Sydney Harbour Estuary sediments was reported by Birch et al., in 2007^12^. The analyses revealed that sediments on the eastern side of the bay had a maximum concentration of 6,920,000 pg.g^-1^ (4352 pg WHO-TEQ g^-1^), with more the 50% of the toxicity being attributed to 2,3,7,8-TCDD^12^. These concentrations placed it among the worlds most PCDD/F contaminated sites, surpassed only by Frierfjorden in the Netherlands, which had a reported maximum PCDD/F concentration of 19,444 pg WHO-TEQ g^-1^ in 2007^12^. Furthermore, PCDD/F were detected more than 15 km from Homebush Bay with the same distinct congener profile, revealing that the Homebush Bay is the contaminant source for the entire estuary.

To date no evidence of PCDD/F degradation has been reported for Sydney Harbour sediments. In anaerobic environments such as those found in marine and estuarine sediments, organohalide respiring bacteria (ORB) can reductively dechlorinate aliphatic and aromatic organohalides by using them as respiratory terminal electron acceptors^13^. Obligate and facultative ORB hail from diverse phyla including: Chloroflexi, Firmicutes and Proteobacteria^14^. However, only *Dehalococcoides mccartyi* strains (Chloroflexi) have been shown to use PCDD/F as terminal electron acceptors (i.e. strains 195^15^, DCMB5^16^, CBDB1^17^, H1−3-2.001 and KKB3.003^18^). Strain CBDB1 is the only strain shown to dechlorinate 2,3,7,8-TCDD^17^. The genus *Dehalococcoides* belongs to the class *Dehalococcoidia*, along with two other obligate organohalide respiring genera (i.e. *Dehalogenimonas, Dehalobium*^19^). While *Dehalococcoides* strains have been shown to be the sole utilizers of PCDD, *Dehalobium* phylotype*s* have been shown to use “dioxin like” polychlorinated biphenyls (PCBs) particularly in estuarine environments^20-23^. Studies on PCDD/F dechlorination in marine environments are rare, and little is known about the microbial communities associated with PCDD/F in marine and estuarine environments.

In this study we investigated PCDD/F reductive dechlorination in Sydney Harbour Estuary sediments by comparing absolute and relative PCDD/F concentrations at the source area with nine surrounding bays, and with previously published data^12^. Microbial community analysis of the sediments revealed an abundance of uncultured marine Dehalococcoidia phylotypes, and one cultured phylotype closely aligned with the *Dehalobium* genus. Attempts to enrich ORB and associated Rdase enzymes with perchloroethene enriched only the *Dehalobium* phylotype. This enrichment culture was then shown to reductively dechlorinate 2,3,7,8-TCDD and OCDD.

## Materials and methods

### Sediment sampling

Sediment sample cores were collected from nine locations along the Parramatta River. Three samples were take from Homebush Bay and six from surrounding bays spanning a distance of 8.5 Km from the PCDD/F source zone (Site 1) in Homebush Bay to Tarban Creek (Site 9) (**Fig. S1, Table S1**). The sediments were manually cored in water less the 2 m in depth by stabbing a 0.5 m long x 0.02 m wide poly carbonate tube attached two a 3 m aluminum long pole into the sediment. The core samples were immediately capped and sealed with PVC tape. With in 4 hours of sampling the sediments cores were transferred to an anaerobic chamber where there were divided into 25 cm upper and lower fractions, transferred to glass bottles and sealed with rubber stoppers under an atmosphere of nitrogen.

### Sample preparation for PCDD/F quantification

Wet sediment was dried overnight at 70°C, the dried sediment was then ground to a fine powder with a mortar and pestle. 20 g of sample was then transferred to an automatic solvent extraction (ASE) cartridge along with 50 g of diatomaceous earth. The sample was spiked with 10 pg.g^-1^ of an isotopically labelled surrogate compound mixture containing ^13^C labelled surrogates of the 17 congeners analyzed (Cambridge Isotope Laboratories Inc. Part # EDF 5393). The samples were extracted with toluene at 200°C and 2000 psi for 10 min (Dionex ASE 350). The toluene was removed under a stream of nitrogen and the residue was reconstituted in dichloromethane (DCM, 1 ml) and then applied to a 5 cm × 1.5 cm column of Florisil. The PCDD/F were eluted with DCM (50 ml). The DCM was removed under a stream of nitrogen; the PCDD/F residue was reconstituted in a toluene (0.5 ml) and transferred to a 2 ml screw cap glass GC vial.

### PCDD/F analysis and quantification

PCDD/F analysis was performed of an Agilent 7890 gas chromatograph (GC) interfaced to an Agilent 7000C triple quadrupole mass spectrometer. The GC was fitted with DB5 (30 m × 0.32 mm (internal diameter) × 0.25 μm (film thickness) column (Agilent technologies, part # 123-5062). The carrier gas (He) flow rate was 0.963 ml.min^-1^. The oven temperature programme was 70°C (1 min) then ramped to 150 °C (15°C.min^-1^), then ramped to 290°C (10°C.min^-1^) and then held at this temperature for 20 mins. The inlet was operated in splitless mode at 250°C. The injection volume was 1 μl and the transfer line was maintained at 290°C. The MS was operated in multiple reaction-monitoring mode (MRM). The mass transitions for each congener and analogous isotopically labelled surrogate are provided in **Table S2**. PCDD/F were quantified by interpolation of a 5-point calibration curve constructed by plotting peak area versus concentration of each congener or surrogate. The concentrations ranged from 0.1 to 1000 ng.mL^-1^ for each congener and 0.1 to 100 ng/ml for surrogates (Cambridge Isotope Labs Inc. Part# EDF-5524) (**Table S3**). The congener of interest was then multiplied by the recovery ratio of its analogous labelled surrogate.

The concentration of each of the 17 congeners was converted to pg WHO-TEQ g^-1^ by multiplying it by its TEF (**Table S4**).

### DNA extraction and 16S rRNA Illumina sequencing

DNA was extracted from sediment using a standard phenol chloroform extraction method. To wet sediment (5 g) in a screw cap polypropylene tube was added 6 mL of lysis buffer (40 mM EDTA, 50 mM Tris-HCl, 0.75 M sucrose, pH 8.0) and 200 μl of lysozyme (125 mg.ml^-1^). The tubes were incubated at 37°C for 1 h after which 4 μl of RNAse A (100 mg.ml^-1^), 5 μl of Proteinase K (20 mg.mL^-1^) and 200 μl of SDS were added to each tube at incubated then incubated at 55°C for 3 h. The tubes were frozen and then thawed the following day at which time 3 ml of aqueous phase was processed by extraction with phenol-chloroform-isoamyl alcohol. DNA was precipitated with isopropanol, washed twice with ice cold 80% ethanol and finally dissolved in 30 μl of DNA-free molecular grade water. Regions of 16S rDNA gene were amplified by PCR from extracted DNA with the Q5 high-fidelity DNA polymerase (New England BioLabs) using the universal primers 926F. 5⍰-TCGTCGGCAGCGTCAGATGTGTATAAGAGACAG - [AAA CTYAAAKGAATTGRCGG]-3⍰) and 1392R (5⍰GTCTCGTGGGCTCGGAGATGTGTATAAGAGACAG -[ACG GGC GGT GTG TRC-3⍰) targeting bacteria and archaea^24^. The samples were sequenced on an Illumina MiSeq Sequencer (Illumina, USA) using V3 chemistry at the Next Generation Sequencing Facility at Western Sydney University’s Hawkesbury Institute for the Environment (Sydney, Australia). 16S rRNA gene amplicon sequences were analyzed with QIIME2-2021.8 utilizing the dada2 pipeline ^25,26^. Sequencing quality was first visualized with FastQC (www.bioinformatics.babraham.ac.uk) resulting in forwards and reverse reads being trimmed to 220 base pairs. Forward and reverse sequences that passed the default quality control were merged and non-overlapping sequences were discarded. Chimeras were analyzed and removed via the consensus method within the dada2 pipeline. Remaining sequences had taxonomy assigned with the RDP classifier ^27^ using the Silva 132 Qiime release database.

### Perchloroethene enrichment culture preparation

To enrich ORB in the harbour sediments, sediments from the most contaminated Site (Site 1, Lower 50 cm, Figure S1A) were used as the inoculum source. The PCE enrichment was prepared using filter-sterilised seawater amended with mineral salts (g/L) [NaCl (1.0), MgCl_2_·H_2_O (0.5), KH_2_PO_4_ (0.2), NH_4_Cl (0.3), KCl (0.3), CaCl_2_·2H_2_O (0.015)], trace elements^28^ and sodium acetate (30 mM). After deoxygenating by nitrogen sparging for 45 min, medium (100 ml) was dispensed into a 120 ml serum flasks containing sediment (1 g). The flasks were sealed with Teflon coated rubber septa and aluminum crimps. The headspace was flushed with N_2_/CO_2_ (4:1) for 3 min. The medium was supplied with 100x vitamin solution^29^, 10 mM NaHCO_3_, and Na_2_S (0.2 mM). The cultures were amended with neat PCE (0.2 mM) using a 10μl glass syringe. Hydrogen gas (0.5 bar) was supplied as an electron donor. All enrichment cultures were incubated statically at 30°C in the dark.

### Chlorinated ethene quantification

Culture headspace gas (100 μl) was withdraw via the septum with a gas tight syringe and manually injected into an Agilent 7890A GC equipped with a flame ionization detector (FID) and a GS-Q capillary column (30 m × 0.32 μm J&W Scientific). The temperature of the split/splitless inlet and detector were set to 250□. The helium carrier gas flow rate was 3 ml/min. The oven program was initially held at 150□, increased to 250□ at a rate of 30□ per min and held for 2 min. The inlet had a split ratio of 1:10.

### Quantitative polymerase chain reaction (qPCR) procedure

The qPCR mixtures contained 2 μl of DNA template, 5 μl of 2×SsoFastTM EvaGreen Supermix, 0.1 μl each of forward and reverse primers, 0.1 μl of Bovine Serum Albumin (BSA) solution and 2.7 μl of DNA-free Molecular grade water. Universal bacterial primers Eub1048F (5’-GTGSTGCAYGGYTGTCGTCA-3’) and Eub1195R (5’-ACGTCRTCCMCACCTTCCTC-3’) were used to quantify the total bacteria concentration^30^. Thermo cycling comprised of 95°C for 3 min, followed by 40 cycles of denaturation at 95°C for 20 seconds and annealing at 62°C for 50 seconds. Following amplification, melting curve analysis was conducted with increments at 0.5°C per 10 seconds from 60 to 99°C. *Dehalococcoides* primers Dehalo505F (5’-GGCGTAAAGTGAGCGTAG-3’) and Dehalo686R (5’-GACAACCTAGAAAACCGC-3’) were used to quantify *Dehalococcoides* spp^31^. Thermo cycling comprised of 98°C for 3 min, followed by 45 cycles of denaturation at 95°C for 30 seconds and annealing at 58°C for 50 seconds. Following amplification, melting curve analysis was conducted with increments at 0.5°C per 5 seconds from 55 to 95°C.

### Sediment microcosms on TCDD and OCDD transformation

TCDD and OCDD were adsorbed onto dried sediments for use as stocks. TCDD and OCDD were dissolved in hexane (50 ml) and mixed with 20 g of dried sediment with the lowest PCDD/F concentration (Site 9). The hexane was evaporated under a stream of nitrogen resulting in 2,3,7,8-TCDD and OCDD fortified sediment with 3.0 μg.g^-1^-dw and 54 μg.g^-1^-dw, respectively.

To test for 2,3,7,8-TCDD and OCDD biotransformation, PCE/sediment enrichment microcosms were purged with N_2_ for 30 min to remove all chloroethenes, and 10% (v/v) inoculated into an anaerobic artificial seawater medium (2% NaCl) and spiked with 1 g of 2,3,7,8-TCDD or OCDD stock dry-sediments. H_2_ (0.5 bar) and sodium acetate (30 mM) were supplied as electron donor and organic carbon source. The H_2_ pressure was returned to 0.5 bar every month. All enrichment cultures were incubated statistically at 30°C in the dark.

### Statistical analysis

Statistical analysis was performed using Prism 9 for MacOS (version 9.1.2). Student’s t-test were two-tallied and unpaired and equal variance was not assumed (unless otherwise stated). Principal component analysis was performed as previously described for PCDD/F in Sydney Harbour Estuary^12^. Absolute PCDD/F concentrations were normalized by dividing by the most abundant congener (either OCDD of OCDF, note Dioxins and furans were treated separately). PCA was then performed on the log_10_ values of the normalized congener concentrations.

## Results and discussion

### PCDD/F concentrations in Sydney Harbour Estuary sediments

The PCDD/F source area adjacent to the former on the eastern side of Homebush Bay was the most contaminated with 1971 × 10^3^ and 4556 × 10^3^ pg.g^-1^ (2026 and 5207 pg WHO-TEQ g^-1^) in upper and lower samples respectively (**Table S5A-D** and **Fig. S2A**). The two other Homebush Bay samples (Sites 2 and 3) had similar concentrations to each other in both upper and lower fractions, with the mean being 540 ± 69 pg WHO-TEQ g^-1^ (n = 4). (**Fig. S2B**). A comparison of mean upper and lower concentrations in the surrounding bays was made with Homebush Bay Sites 2 and 3 (the source was excluded). The comparison showed that there was no significant difference in the total PCDD/F concentrations up to 4.5 km away (*p* > 0.05). However, at 8 km the PCDD/F concentrations were significantly lower (**Fig. S2B**).

In all 18 samples the PCDDs where in greater abundance than PCDFs. When averaged across all sampling sites PCDDs were 18.2 ± 4.0 and 21.3 ± 11.5 times greater than PCDFs in upper and lower sediments respectively (**Table S5A & B**). The perchlorinated congeners OCDD or OCDF were consistently the most abundant within their chemical class (i.e. PCDD or PCDF) across all nine locations at both depths. OCDD accounted for 96.3 ± 8.4% (n = 18) of the PCDDs and OCDF accounted for 94.3 ± 7.7% (n = 18) of PCDFs. 2,3,7,8-TCDD accounted for 0.02 ± 0.006% of the PCDD/F mass and 15.9 ± 8.4% (n = 18) of the toxicity (**Table S5 A-D**).

### Principal component analysis of PCDD/F congener profiles

Principal component analysis (PCA) of normalized congener profiles previously showed a tight clustering of samples from Homebush Bay with surrounding areas^12^. From this correlation the authors concluded that the PCDD/F contamination at Homebush Bay was the source of contamination for surrounding areas of the estuary extending 5 km to the west and 15 km to the east of the source. Similarly, we performed PCA on log_10_ of normalized congener concentrations at all sample locations. Normalized congener concentrations were achieved by dividing by the most abundant congener (either OCDD of OCDF, note Dioxins and furans were treated separately). The Homebush Bay congener profiles from 2007 were treated in the same way and included for comparison (**Fig 1A**). As expected the Homebush Bay congener profiles from 2007 clustered together. In contrast the tight clustering of samples was no longer evident in congener profiles determined in the present study. The source area is clearly different to Sites 2 and 3 in Homebush Bay (that cluster together) and surrounding bays (Sites 4-9). These results suggest that significant changes in the congener profiles have occurred in the 14 years between studies particularly in Homebush Bay.

**Figure 1.**
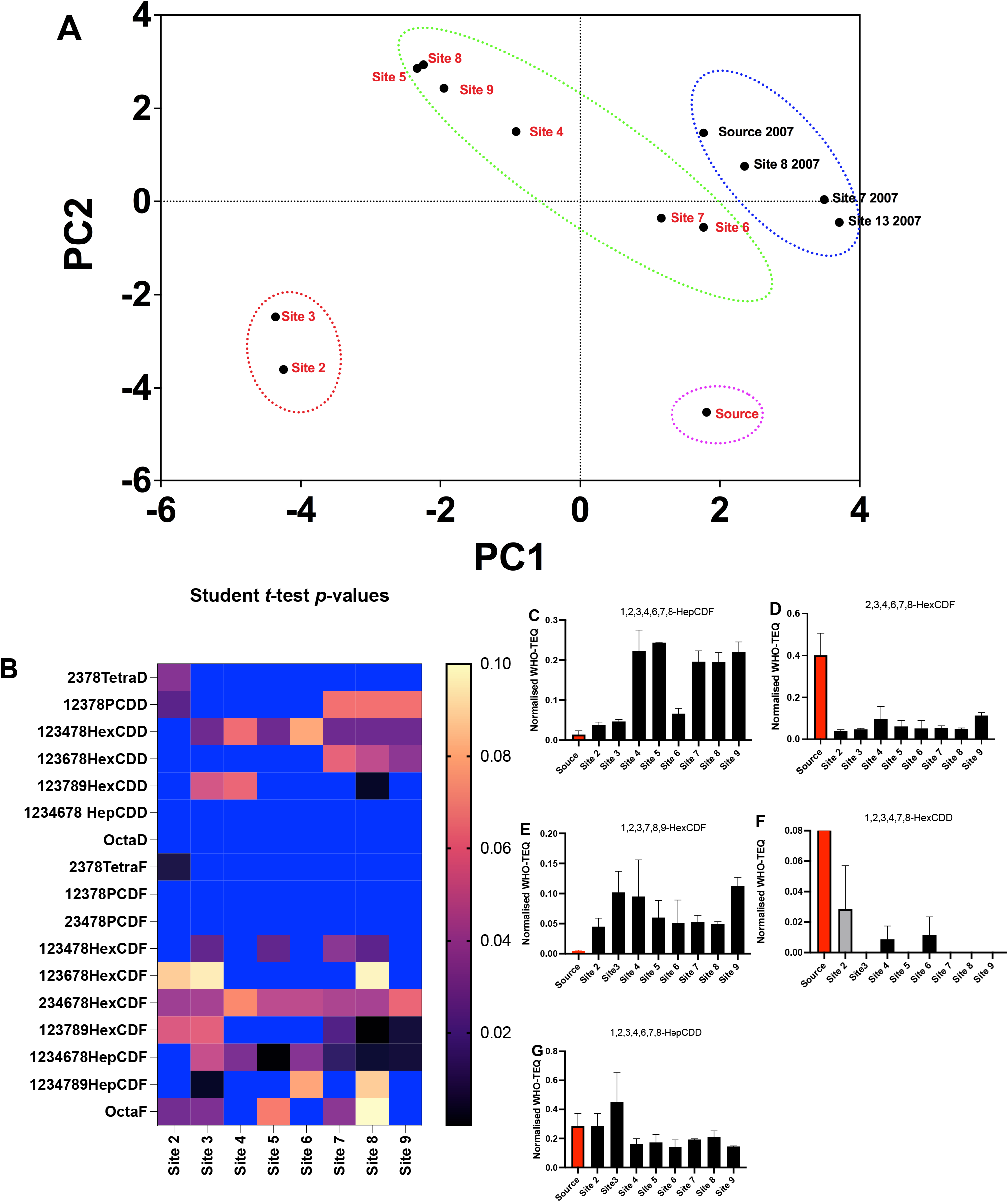
**Panel A**. PCA plot of congener profiles at Homebush Bay and surrounding Bays. Each congener concentration was normalized to the total congener concentration and then transformed (log_10_). Data from 2007 were included for comparison (circled in blue). Homebush Bay 2017 samples (circled in red), 2017 Surrounding Bays (circled in green), 2017 source zone (circled in pink). **Panel B**. t-test comparison of mean (upper and lower) normalized congener concentrations at the source zone with the other eight locations. (dark blue indicates a p-value > 0.1). **Panels C-G** shows the normalized congener concentrations that are significantly different to those at the source

To elucidate the significant differences in congener profiles at the source compared to other sites, a pair wise comparison was made in the relative abundance of each congener at the source with the corresponding congener at each of the sampling sites (**Fig. 1B**). The pairwise *t-*test comparisons revealed significant (*p* < 0.05) differences in PCDFs, 1,2,3,4,6,7,8-HepCDF, 1,2,3,7,8,9- and 2,3,4,6,7-HexCDF (**Fig 2C**). A closer inspection of these congeners revealed a lower relative abundance of HepCDF, accompanied by elevated relative abundance of 2,3,4,6,7,8-hexaCDF at the source compared with the other locations, suggesting that reductive dechlorination of the hepta congener has occurred at the source (**Fig 2B & C**). The proportion of 1,2,3,7,8,9-hexaCDF was also significantly lower at the source relative to all other locations (**Fig. 2D**). However, there was no significance difference in the proportion of penta congeners, suggesting that 1,2,3,7,8,9-hexaCDF was transformed to a penta congener that was not amongst the suite of compounds analyzed. Regarding PCDDs 1,2,3,4,7,8-HexCDD was significantly higher in relative abundance at the source than at all sites, a potential precursor to 1,2,3,4,7,8-HexCDD, (i.e. 1,2,3,4,6,7,8-heptaCDD was not significantly different in relative abundance across the nine sites (**Fig 2F**).

**Figure 2.**
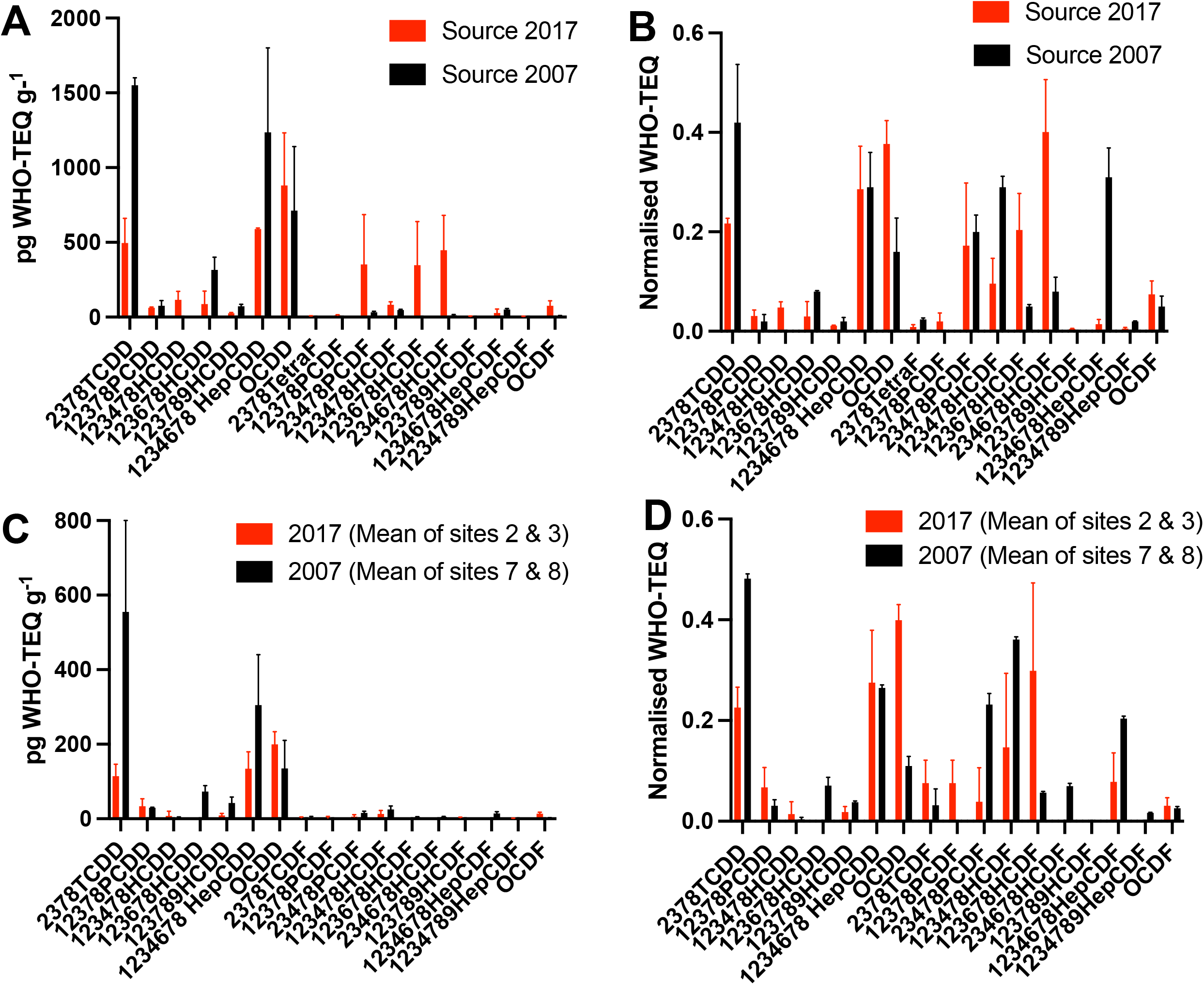
Homebush Bay PCDD/F congener concentrations and normalized concentrations in 2007 (historical^12^) and 2017 (present study). **Panel A-B** Source area on the eastern side the bay. **Panels C-D** sample locations outside the source area (Precise locations are shown in Figure S3).

### Comparison of Homebush Bay congener profiles with historical data indicates 2,3,7,8-TCDD degradation

The Homebush Bay sediment-sample locations in the current study were taken from similar areas to those in 2007^12^ (**Fig S3**). Samples were grouped into those at the source area and those outside the source area for comparison of congener TEQs and profiles (**Fig 3A & C**). In both cases the most significant congener in 2007 in terms of toxicity was 2,3,7,8-TCDD (**Fig. 3A&C**). However in the present study the 2,3,7,8-TCDD was significantly lower than historical levels both inside (*p* = 0.013) and outside (*p* = 0.015) the source area. The reduced abundance of 2,3,7,8-TCDD was also evident in the normalized congener TEQ profile demonstrating that 2,37,8-TCDD had diminished in abundance relative to all other congeners as well as relative to historical values (**Fig 3B & D**).

**Figure 3.**
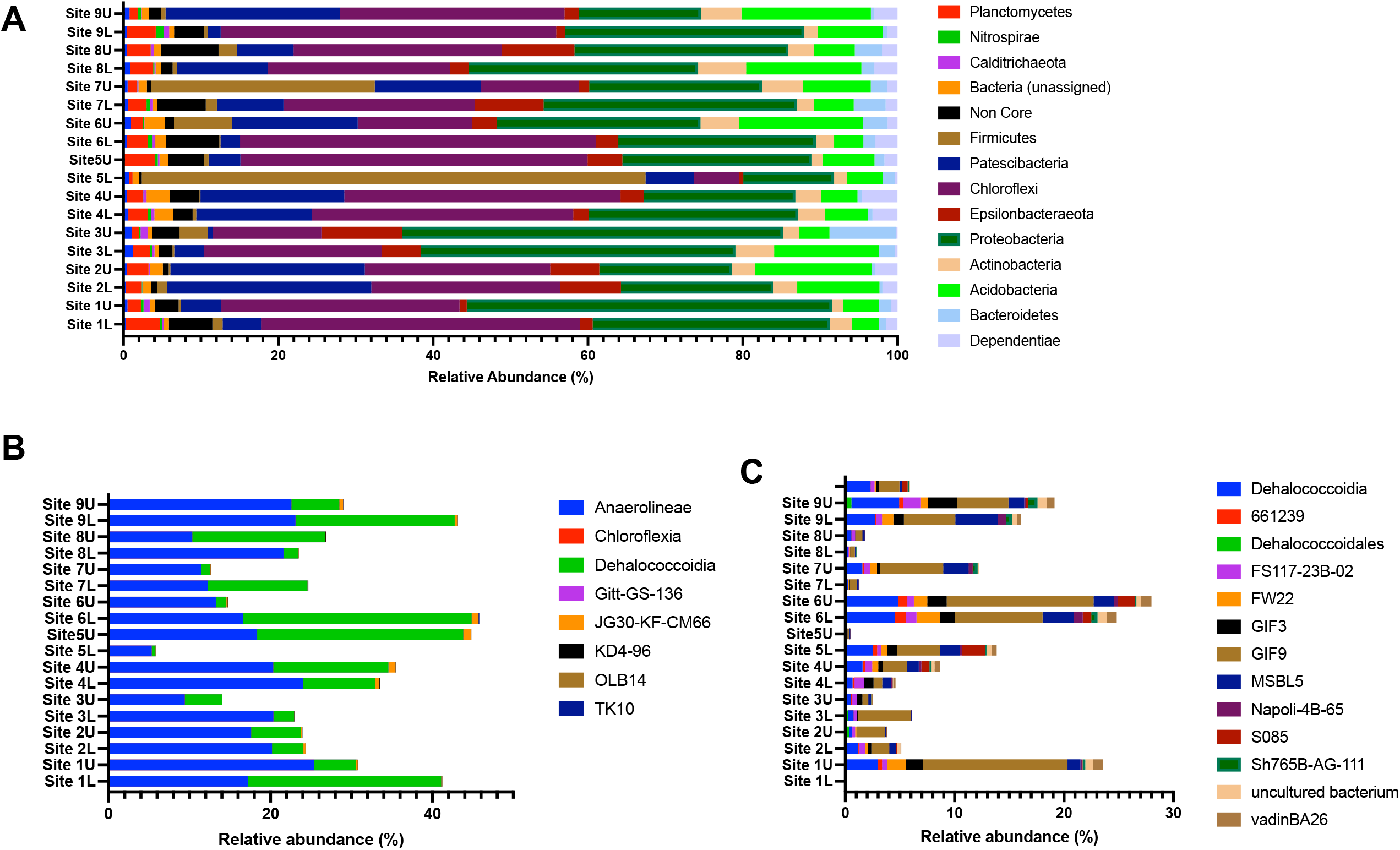
**Panel A**. Relative abundance of the core bacterial phyla in Sydney Harbour sediments at the level of Phylum. **Panel B**. Class within the Chloroflexi phyla. **Panel C**. Order within the Dehalococcoidia class (**C**).

Analysis of the sediment extracts for 2,7,8-triCDD, 2,7-/2,8-DiCDD, chloroCDD revealed all but 2,7,8-TriCDD to be present suggesting that reductive dechlorination could be a mechanism of 2,3,7,8-TCDD transformation (**Fig S4**). However, as there is no prior data for lesser-chlorinated PCDDs in the harbour sediments it is not known if they were part of the initial pollutant mix or if they are evidence of *in situ* 2,3,7,8-TCDD reductive dechlorination. Nevertheless, this study provides baseline data for future comparison.

### 16S amplicon sequencing reveals potential PCDD/F respiring bacteria

The microbial community analysis in this study was performed primarily to identify ORB that have the potential to reductively dechlorinate PCDDs, and specifically 2,3,7,8-TCDD (i.e. *Dehalococcoides* mccartyi strains). DNA extracted directly from the sediment samples was analyzed by 16S rRNA gene amplicon sequencing. The number of sequences retrieved ranged between 49158 to 115942 (mean = 77216 ± 22914 (n = 18). The community consisted of 15 phyla with a relative abundance above 0.5% of which 12 were represented in all samples (i.e. the microbial core community) (Fig 3A). The core community represented >94% of identified phyla; the non-core community ranged between 0.7 and 6%.

Chloroflexi was the most abundant phyla accounting for 28.1 ± 11.3% of the community across all samples (n=18) (**Fig 3B**). The class Dehalococcoidia in the Chloroflexi phylum accounted for 10.9 ± 9.30 % of the total community second to Anaerolineae (16.8± 5.37%). Previously, 16S rRNA phylogenetic analysis of orders within Dehalococcoidia have shown them to assemble in three clusters (i.e. Marine I, Marine II, and Terrestrial I)^32^. All cultivated phylotypes belong to Terrestrial Cluster I, and are characterised by their ability to conserve energy via respiration of organohalides^32^. The physiological role of Dehalococcoidia in marine sediments is unclear primarily due to a lack of cultivated phylotypes. However, analysis of metagenome and single cell assembled genomes (MAGS and SAGS) of marine Dehalococcoidia have revealed a general absence of reductive dehalogenase genes suggesting that they conserve energy though means other than organohalide respiration^33, 34^. In the present study Dehalococcoidia affiliates were denominated as uncultivated lineages with the order GIF9 (Marine Cluster II) being the most abundant across the 18 samples (38.2 ± 15.8% of Dehalococcoidia) (Fig 3C). GIF9 has been shown to possess only a partial reductive dehalogenase gene and therefore it seems unlikely that it is participating in PCDD/F dechlorination^32^. *Dehalobium* in the Dehalococcoidales order was the only cultivated lineage identified at the genus level and accounted for a small proportion of the Dehalococcoidia class (0.33 ± 0.51% (n = 18)). *Dehalobium* species have been shown to use PCE and PCBs as terminal electron acceptors in marine environments^20-22^. However, there are no reports of *Dehalobium* species dechlorinating PCDD/F.

### Enrichment of *Dehalobium* spp. using PCE as electron acceptor

Given the lack of clarity regarding ORB within the sediments it was decided to attempt to enrich the ORB population using PCE. PCE was chosen because *Dehalococcoides* mccartyi strain CBDB1 is capable of using both PCE and 2,3,7,8-TCDD as its terminal electron acceptor^17, 35^. However, PCE with an aqueous solubility of 150 mg.L^-1^ has far greater bioavailability than 2,3,7,8-TCDD with an aqueous solubility of 0.2 μg.L^-1 (36)^.

The enrichment cultures were prepared with sediment from source zone (lower fraction) with filter-sterilized seawater to replicate *in situ* conditions. The first PCE (0.2 mM) pulse was dechlorinated to TCE (0.10 mM) and *cis*-DCE (0.06 mM). Additional PCE (0.2 to 0.4 mM) was subsequently supplied. After 85 days the best performing microcosm had dechlorinated 2.60 mM PCE to TCE (1.41 mM) and cis- and trans-DCE (0.74 and 0.44 mM, respectively) (**Fig 4A**). No further dechlorination to vinyl chloride or ethene was observed. Sulfate reduction was also observed and was completely depleted from the seawater medium (data not shown). Neither PCE or sulfate reduction occurred in sterilized controls.

**Figure 4.**
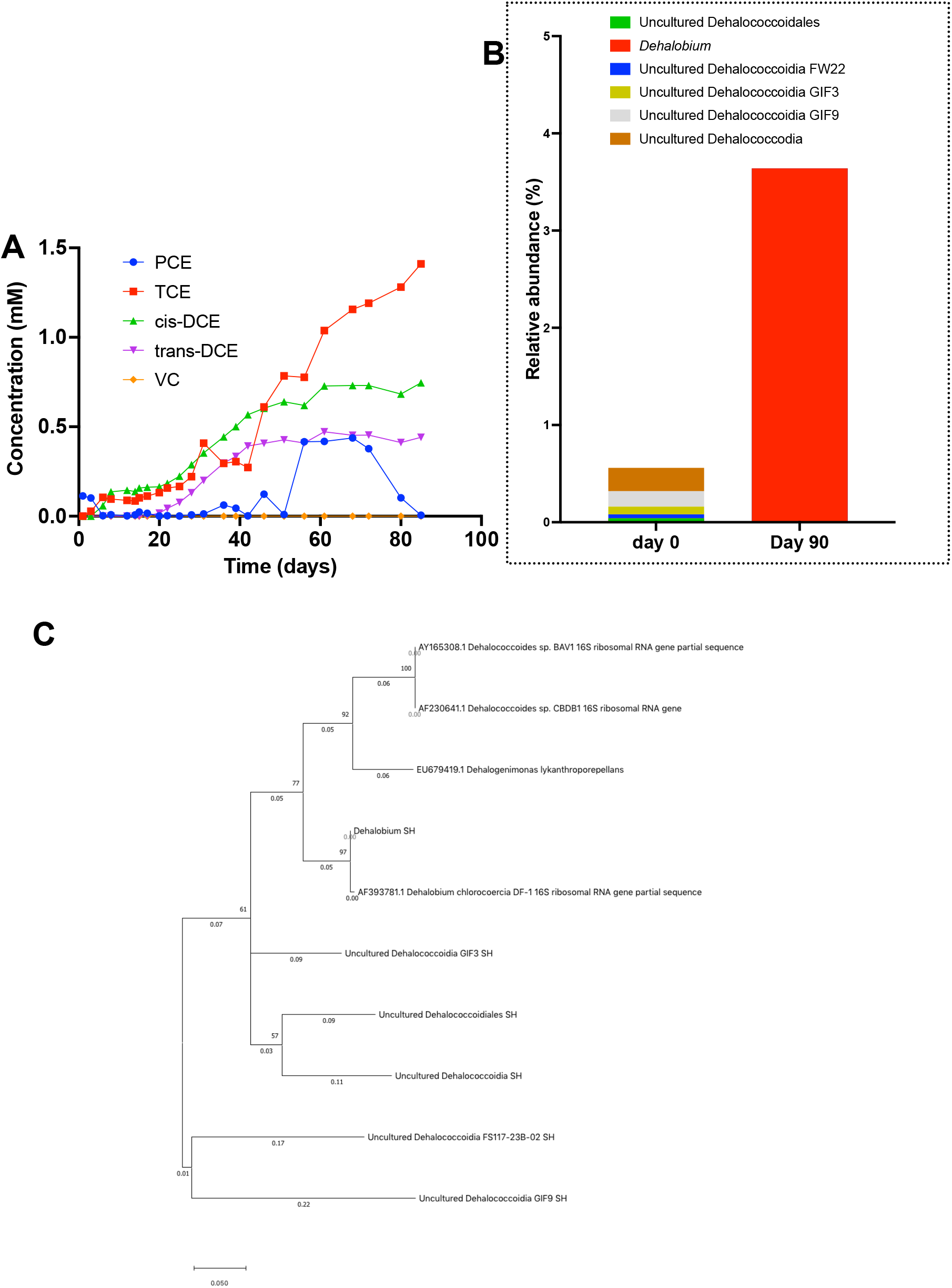
**Panel A**. PCE dechlorination by a sediment enrichment culture in seawater supplied with hydrogen and acetate as electron donor and carbon source respectively. **Panel B** 16S rRNA community analysis of the PCE enrichment culture at time zero and day 90. **Panel C** Maximum likely-hood Phylogenetic tree of Dehalococcoidia partial 16S rRNA gene sequences acquired in this study from Sydney Harbour Estuary sediments (labelled SH) relative to cultivated phylotypes in the Dehalococcoidia class. Sequence alignment, modeling and construction were performed with MEGA (version 10.2.6). Numbers next to the nodes represent the percentage of branch support based on bootstrap re-sampling (500x).

The PCE-enrichment culture was analyzed by Illumina 16S rRNA gene amplicon sequencing at day 0 and 90 (**Fig 4B**). After 90 days, the only ORB was identified in the community was *Dehalobium* from the Dehalococcoidia class, (3.6 % of the community) (**Fig 4B**). No other Dehalococcoidia lineages that were present at day zero were detectable at day 90. The production of both *cis-* and *trans-*DCE in a ratio of 1.5 is consistent with previous reports for *Dehalobium* lineages when respiring PCE^37^. Blastn alignment of the 266 base-pair representative sequences aligned most closely with uncultured Dehalococcoidia VLD-1 (98.95% identity), which is closely related to *Dehalobium* chlorocoercia DF-1 (**Fig 4C**). A large increase in the relative abundance of sulfate reducing bacteria including *Desulfovibrio* (52%) and the Desulfobulbaceae family (15%) was observed. Their proliferation is consistent with the observed removal of sulfate from the seawater medium.

### TCDD and OCDD dechlorinating activity in ORB enrichment culture

The PCE enrichment culture was applied in anaerobic cultures containing 2,3,7,8-TCDD and OCDD as electron acceptors (i.e. the congener with the highest TEQ and the most abundant by mass respectively) and hydrogen as the electron donor. A separate set of cultures was prepared that included the biosurfactant lecithin, that has been shown to increase the bioavailability of lipophilic organic pollutants such as PCDD/F and PCBs^38, 39^.

The enrichment cultures resulted in the reductive dechlorination of both congeners after a lag of five-months. In 2,3,7,8-TCDD amended microcosms, 2,3,7-TriCDD was detectable in the lecithin treatments after 5-months, but not in the lecithin free control. After 14-months 2,3,7,-TriCDD was detected in both lecithin and lecithin free treatments with no statistical difference between the two (**Fig 5A**). After 28 months of observation there was more 2,3,7-TriCDD produced in the lecithin amended microcosms (0.49 ± 0.3 nmoles.g^-1^ vs. 0.10 ± 0.07 nmoles.g^-1^ (*p* = 0.093). The quantity of 2,3,7-TriCDD produced represented 5.4 ± 1.0% and 1.1 ± 0.77 mole % of the initial amount of 2,3,7,8-TCDD added to the lecithin and lecithin free microcosms respectively.

**Figure 5.**
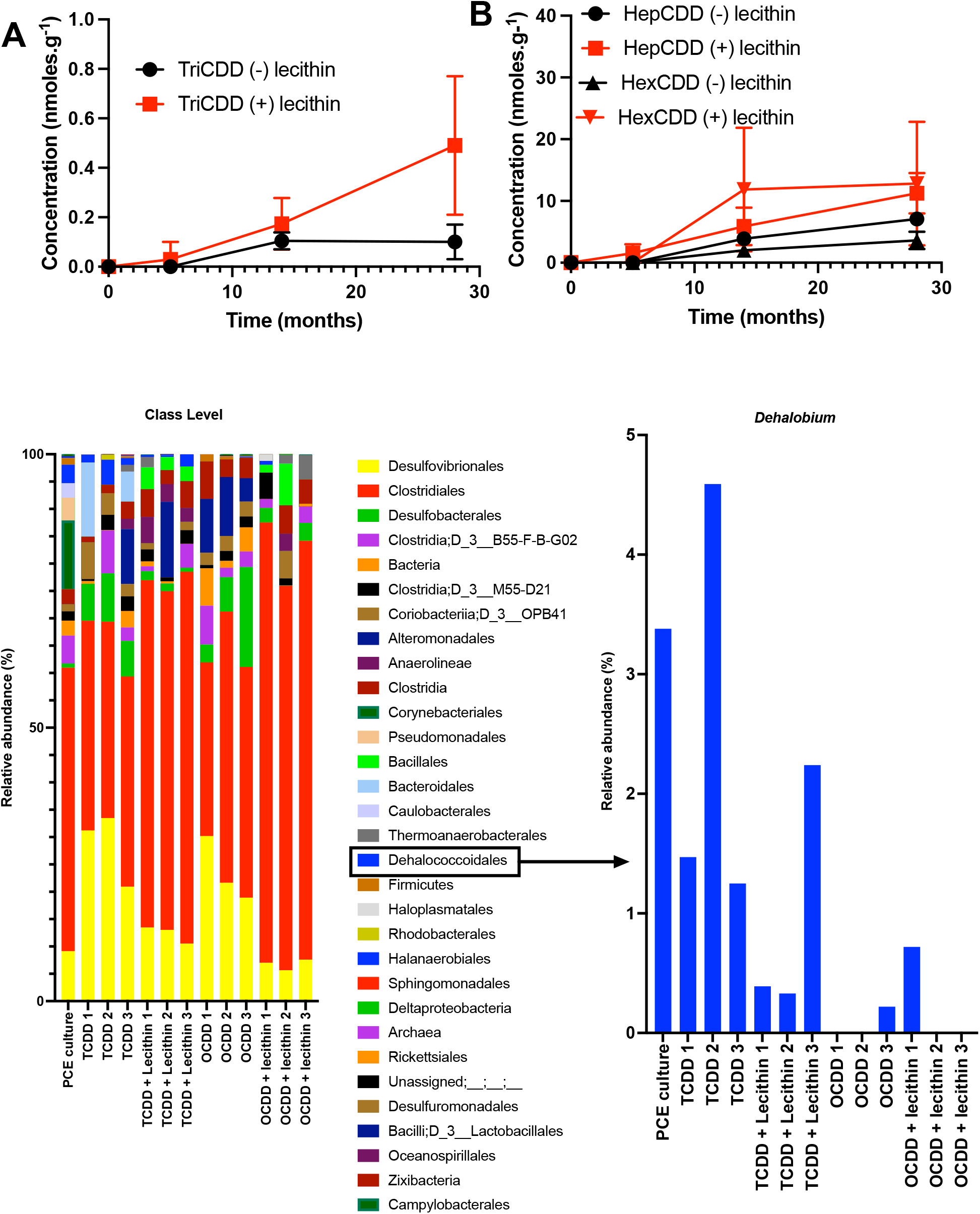
The production of 2,3,7-TriCDD from 2,3,7,8-TCDD (**Panel A**) and 1,2,3,4,6,7,8-HepCDD and 1,2,3,4,7,8-HexCDD from OCDD (**Panel B**) in anaerobic seawater +/-lecithin microcosms inoculated with the PCE/*Dehalobium* enrichment culture. Error bars represent standard deviation (n = 3). **Panel C** 16S rRNA community analysis of the 2,3,7,8-TCDD and OCDD microcosms inoculated with PCE/*Dehalobium* enrichment culture after 14 months of incubation.

OCDD was dechlorinated to 1,2,3,4,6,7,8-HepCDD and 1,2,3,4,7,8-HexCDD (**Fig. 5B**). HepCDD was detectable after five months where lecithin was employed but not in the lecithin free control. After 28 months of observation the sum of dechlorination products in lecithin-amended microcosms (24.1 ± 10.1 nmol.g^-1^) was significantly higher at the 90% confidence interval than those without lecithin (10.7 ± 1.59 nmol.g^-1^) (*p* = 0.089). No dechlorination was observed in sterile controls, suggesting that the transformation was dependent on microbial activity.

16S rRNA sequencing of DNA from the cultures after 14 months showed that no other member of Dehalococcoidia class besides *Dehalobium* were detectable during 2,3,7,8-TCDD and OCDD dechlorination. However, during OCDD dechlorination the *Dehalobium* population diminished significantly to below detection in four of six microcosms, whereas in 2,3,7,8-TCDD microcosm the *Dehalobium* population ranged from 0.33 – 4.59% of the microbial population (Fig 5C). This suggests that OCDD dechlorination may not be linked to *Dehalobium*. The mechanism of dechlorination of OCDD dechlorination could be an abiotic process via reduced cobalamins in the medium. A previous study has shown that reduced cyanocobalamin can catalyze the reductive dechlorination of OCDD to TCDD but not beyond^40^.

Cultures amended with lecithin had increased relative abundance of Clostridiales, known to ferment a range of organic compounds under anaerobic conditions^41^. The relative abundance of Clostridiales in all lecithin-amended cultures accounted for 70 ± 6.6% (n = 6) of the microbial population compared with 39.4 ± 5.5% (n = 6) of the population in unamended cultures. This finding suggests that lecithin is being degraded by members of the Clostridiales and hence is utility as long-term biosurfactant to solubilize PCDD/F is doubtful.

## Conclusion

In the study we have insight into the natural attenuation of PCDD/F congeners in the sediments in Sydney Harbour Estuary. The most significant transformation observed was a 50% decrease in the most toxic and concerning congener 2,3,7,8-TCDD accompanied by the presence of lesser-chlorinated dioxins in the sediments. Taken together these findings suggest that reductive dehalogenation is at least one mechanism by which 2,3,7,8-TCDD is being detoxified. Microbial community analysis of the sediments revealed an abundance of lineages from the Dehalococcoidia class. However, the majority of these lineages were associated with physiologically uncharacterized marine phylotypes whose ability to use organohalides as terminal electron acceptor is in despute. Nevertheless an ORB from Dehalococcoidia was enriched using PCE as the electron acceptor that phylogenetically aligned with the *Dehalobium* genus. The *Dehalobium* enrichment culture was then shown to reductively dechlorinate 2,3,7,8-TCDD and OCDD. Collectively, these findings show promise for the bioremediation of marine sediments contaminated with PCDD/F. Future work will entail defining the repertoire of PCDD/F congeners and other organohalides that this culture can reductively dechlorinate and genetically and functionally characterizing the reductive dehalogenase(s) that catalyze these reactions. PCR primers based RDase gene sequences coupled with the 16S rRNA phylogenetic marker genes will be valuable diagnostic tool available to remediation practitioners to determine the *in situ* capability for PCDD/F attenuation.

