## Supplementary data for "*Dehalobium* species implicated in 2,3,7,8-tetrachloro-*p*-dioxin dechlorination in the contaminated sediments of Sydney Harbour Estuary"

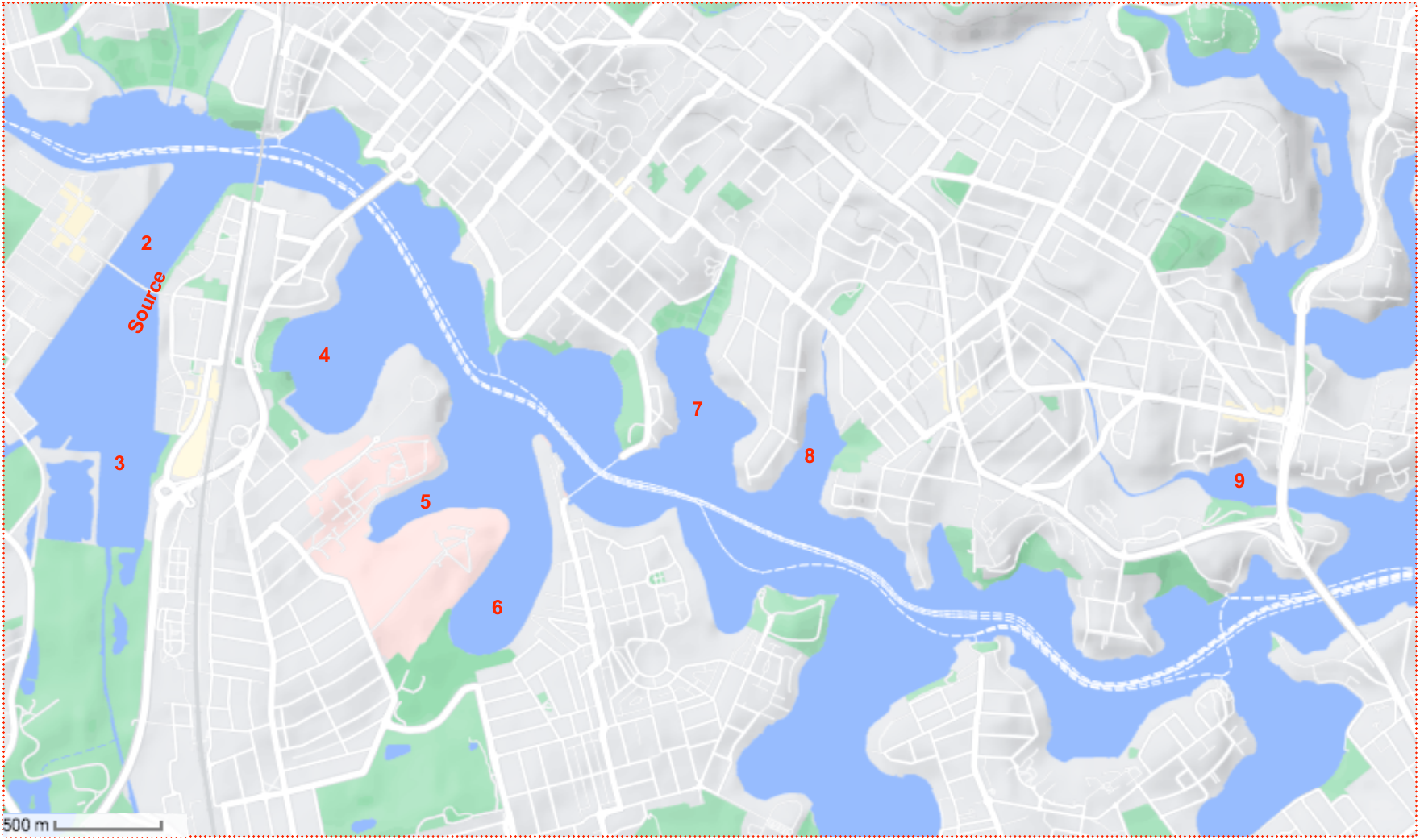

**Figure S1:** Sediment sampling location in Homebush Bay (Source, Sites 2 and 3) and surrounding bays heading east along the Parramatta River towards Tarban Creek (9) (co-ordinates for each sample location are provided in table S2).

**Table S1:** Details of sampling sites locations shown in Figure S1.

| **Site number** | **Latitude** | **Longitude** | **Distance from source** | **Name of location** |
| --- | --- | --- | --- | --- |
| Source | -33.8269 | 151.0830 | 0 | Homebush Bay |
| 2 | -33.8249 | 151.0824 | 239 | Homebush Bay |
| 3 | -33.8361 | 151.0803 | 1000 | Homebush Bay |
| 4 | -33.8319 | 151.0940 | 2250 | Brays Bay |
| 5 | -33.8406 | 151.1038 | 3160 | Majors Bay |
| 6 | -33.8377 | 151.0971 | 3370 | Yaralla Bay |
| 7 | -33.8308 | 151.1121 | 4130 | Morrison’s Bay |
| 8 | -33.8337 | 151.1195 | 4470 | Glades Bay |
| 9 | -33.8367 | 151.1416 | 8279 | Tarban Greek |

**Table S2:** Mass spectrum Multiple Reaction Monitoring Mode (MRM) parameters. Ion polarity was positive in all cases

| **Compound Name** | **RT** | **Quantifier** | | | **Qualifier** | | |
| --- | --- | --- | --- | --- | --- | --- | --- |
|  |  | **Precursor ion** | **Product ion** | **CE (V)** | **Precursor ion** | **Product ion** | **CE (V)** |
| Dibenzo-p-dioxin (DD) | 11.8 | 184.0 | 128.1 | 25 | 184.0 | 102.1 | 25 |
| MCDD | 13.2 | 217.9 | 155.0 | 25 | 217.9 | 127.1 | 25 |
| 2,7-DCDD | 14.8 | 251.9 | 189.0 | 25 | 251.9 | 217.0 | 25 |
| 2,3-DCDD | 14.9 | 251.9 | 189.0 | 25 | 251.9 | 217.0 | 25 |
| 2,3,7-TriCDD | 17.1 | 285.8 | 223.0 | 25 | 285.8 | 251.1 | 25 |
| 2,3,7,8-TCDD | 20.1 | 321.8 | 259.0 | 24 | 321.8 | 257.0 | 24 |
| 1,2,3,7,8-PCDD | 22.6 | 353.9 | 292.9 | 25 | 353.9 | 290.9 | 25 |
| 1,2,3,4,7,8-HexCDD | 27.5 | 389.8 | 326.9 | 25 | 391.8 | 328.8 | 25 |
| 1,2,3,6,7,8-HexCDD | 27.6 | 389.8 | 326.9 | 25 | 391.8 | 328.8 | 25 |
| 1,2,3,7,8,9-HexCDD | 28.0 | 389.8 | 326.9 | 25 | 391.8 | 328.8 | 25 |
| 1,2,3,4,6,7,8-HepCDD | 30.5 | 423.8 | 360.8 | 25 | 425.8 | 362.8 | 25 |
| OCDD | 37.8 | 457.7 | 394.8 | 26 | 459.5 | 396.8 | 26 |
| 2,3,7,8-TCDF | 19.9 | 303.9 | 240.9 | 33 | 305.9 | 242.9 | 33 |
| 1,2,3,7,8-PCDD | 22.9 | 339.9 | 276.9 | 35 | 337.9 | 274.9 | 35 |
| 2,3,4,7,8-PCDF | 23.5 | 339.9 | 276.9 | 35 | 337.9 | 274.9 | 35 |
| 1,2,3,4,7,8-HexCDF | 26.6 | 373.8 | 310.9 | 35 | 375.8 | 312.9 | 35 |
| 1,2,3,6,7,8-HexCDF | 26.7 | 373.8 | 310.9 | 35 | 375.8 | 312.9 | 35 |
| 2,3,4,6,7,8-HexCDF | 27.3 | 373.8 | 310.9 | 35 | 375.8 | 312.9 | 35 |
| 1,2,3,7,8,9-HexCDF | 28.3 | 373.8 | 310.9 | 36 | 375.8 | 312.9 | 36 |
| 1,2,3,4,6,7,8-HepCDF | 32.4 | 407.8 | 344.8 | 36 | 409.8 | 346.8 | 36 |
| 1,2,3,4,7,8,9-HepCDF | 30.7 | 407.8 | 344.8 | 35 | 409.8 | 346.8 | 36 |
| OCDF | 38.9 | 443.7 | 380.6 | 35 | 441.7 | 378.8 | 35 |

**Table S3** Mass spectrum Multiple Reaction Monitoring Mode (MRM) parameters for isotopically labelled internal standards. Ion polarity was positive in all cases

| **Congener** | **TEF** |
| --- | --- |
| 2,3,7,8-TCDD | 1 |
| 1,2,3,7,8-PCDD | 1 |
| 1,2,3,4,7,8-HexCDD | 0.1 |
| 1,2,3,6,7,8-HexCDD | 0.1 |
| 1,2,3,7,8,9-HexCDD | 0.1 |
| 1,2,3,4,6,7,8-HepCDD | 0.01 |
| OCDD | 0.0003 |
| 2,3,7,8-TCDF | 0.1 |
| 1,2,3,7,8-PCDF | 0.03 |
| 2,3,4,7,8-PCDF | 0.3 |
| 1,2,3,4,7,8-HexCDF | 0.1 |
| 1,2,3,6,7,8-HexCDF | 0.1 |
| 2,3,4,6,7,8-HexCDF | 0.1 |
| 1,2,3,7,8,9-HexCDF | 0.1 |
| 1,2,3,4,6,7,8-HepCDF | 0.01 |
| 1,2,3,4,7,8,9-HepCDF | 0.01 |
| OCDF | 0.0003 |

**Table S4:** World Health Organization toxicological equivalence factors (WHO-TEF)

| **Congener** | **Source** | **Site 2** | **Site 3** | **Site 4** | **Site 5** | **Site 6** | **Site 7** | **Site8** | **Site 9** |
| --- | --- | --- | --- | --- | --- | --- | --- | --- | --- |
| 2,3,7,8-TCDD | 330.4 | 142.5 | 78.9 | 95.5 | 27.1 | 110.8 | 94.9 | 102.9 | 30.5 |
| 1,2,3,7,8-PCDD | 68.2 | 45.0 | 0.0 | 44.2 | 41.8 | 62.2 | 0.0 | 0.0 | 0.0 |
| 1,2,3,4,7,8-HexCDD | 583.3 | 300.8 | 0.0 | 0.0 | 0.0 | 0.0 | 0.0 | 0.0 | 0.0 |
| 1,2,3,6,7,8-HexCDD | 0.0 | 0.0 | 0.0 | 466.9 | 309.4 | 588.2 | 400.8 | 420.2 | 285.9 |
| 1,2,3,7,8,9-HexCDD | 199.5 | 0.0 | 93.4 | 101.8 | 59.4 | 142.3 | 0.0 | 77.7 | 61.2 |
| 1,2,3,4,6,7,8-HepCDD | 59617 | 11033.9 | 21217.2 | 9552.6 | 2378.2 | 13534.9 | 5959.0 | 5107.6 | 2082.6 |
| OCDD | 17615178 | 667663.2 | 564353.1 | 628779.8 | 237555.5 | 1100594.8 | 407117.6 | 360315.2 | 178702.4 |
| 2,3,7,8-TCDF | 57.6 | 39.4 | 0.0 | 43.3 | 0.0 | 0.0 | 0.0 | 0.0 | 0.0 |
| 1,2,3,7,8-PCDD | 522.0 | 0.0 | 0.0 | 0.0 | 0.0 | 0.0 | 0.0 | 0.0 | 0.0 |
| 2,3,4,7,8-PCDF | 66.3 | 34.0 | 41.1 | 38.5 | 0.0 | 315.8 | 0.0 | 0.0 | 0.0 |
| 1,2,3,4,7,8-HexCDF | 627.3 | 0.0 | 151.9 | 161.1 | 73.1 | 258.6 | 115.4 | 192.4 | 91.8 |
| 1,2,3,6,7,8-HexCDF | 554.0 | 0.0 | 0.0 | 0.0 | 0.0 | 0.0 | 0.0 | 0.0 | 0.0 |
| 2,3,4,6,7,8-HexCDF | 2160.0 | 0.0 | 0.0 | 0.0 | 0.0 | 0.0 | 0.0 | 0.0 | 0.0 |
| 1,2,3,7,8,9-HexCDF | 25.7 | 26.0 | 31.8 | 0.0 | 0.0 | 0.0 | 0.0 | 0.0 | 0.0 |
| 1,2,3,4,6,7,8-HepCDF | 0.0 | 0.0 | 0.0 | 993.5 | 552.8 | 1215.6 | 700.3 | 650.1 | 396.1 |
| 1,2,3,4,7,8,9-HepCDF | 343.1 | 88.9 | 176.2 | 72.8 | 0.0 | 185.9 | 131.7 | 79.4 | 0.0 |
| OCDF | 144047 | 57714 | 30821.8 | 31745.3 | 12529.5 | 40057.1 | 18466.1 | 16918.7 | 3561.7 |
| **Total** | **1970719** | **737087** | **616965** | **672095** | **253526** | **1157066** | **432985** | **383864** | **185212** |

**Table S5A:** Concentration (pg.g^-1^) of individual congeners in the upper 25cm of the sediment cores

**Table S5B:** Concentration (pg.g^-1^)of individual congeners in the lower 25cm of the sediment cores

| **Congener** | **Source** | **Site 2** | **Site 3** | **Site 4** | **Site 5** | **Site 6** | **Site 7** | **Site 8** | **Site 9** |
| --- | --- | --- | --- | --- | --- | --- | --- | --- | --- |
| 2,3,7,8-TCDD | 661.1 | 142.5 | 87 | 95.5 | 0.0 | 27.1 | 159.6 | 110.8 | 0.0 |
| 1,2,3,7,8-PCDD | 55.8 | 45.0 | 41 | 44.2 | 31.8 | 41.8 | 0.0 | 62.2 | 0.0 |
| 1,2,3,4,7,8-HexCDD | 1721.7 | 300.8 | 0 | 0.0 | 23.0 | 0.0 | 0.0 | 0.0 | 0.0 |
| 1,2,3,6,7,8-HexCDD | 1736.2 | 0.0000 | 0 | 466.9 | 285.6 | 309.4 | 268.8 | 588.2 | 315.5 |
| 1,2,3,7,8,9-HexCDD | 304.4 | 0.0 | 113 | 101.8 | 45.0 | 59.4 | 130.0 | 142.3 | 0.0 |
| 1,2,3,4,6,7,8-HepCDD | 58060.8 | 11033.9 | 10286 | 9552.6 | 1637.3 | 2378.2 | 17847.5 | 13534.9 | 1407.8 |
| OCDD | 4108152.3 | 667663.2 | 578631 | 628779.8 | 160809.4 | 237555.5 | 1350515.3 | 1100594.8 | 181073.9 |
| 2,3,7,8-TCDF | 93.6 | 39.4 | 36 | 43.3 | 0.0 | 0.0 | 0.0 | 0.0 | 0.0 |
| 1,2,3,7,8-PCDD | 259.5 | 0.0 | 0 | 0.0 | 0.0 | 0.0 | 0.0 | 0.0 | 0.0 |
| 2,3,4,7,8-PCDF | 2285.8 | 34.0 | 0 | 38.5 | 0.0 | 0.0 | 0.0 | 315.8 | 0.0 |
| 1,2,3,4,7,8-HexCDF | 1027.5 | 0.0 | 128 | 161.1 | 0.0 | 73.1 | 237.8 | 258.6 | 0.0 |
| 1,2,3,6,7,8-HexCDF | 6383.9 | 0.0 | 0 | 0.0 | 0.0 | 0.0 | 0.0 | 0.0 | 0.0 |
| 2,3,4,6,7,8-HexCDF | 6802.7 | 0.0 | 0 | 0.0 | 0.0 | 0.0 | 0.0 | 0.0 | 0.0 |
| 1,2,3,7,8,9-HexCDF | 72.7 | 26.0 | 53 | 0.0 | 0.0 | 0.0 | 0.0 | 0.0 | 0.0 |
| 1,2,3,4,6,7,8-HepCDF | 5468.1 | 0.0 | 0 | 993.5 | 352.9 | 552.8 | 1538.7 | 1215.6 | 386.7 |
| 1,2,3,4,7,8,9-HepCDF | 542.0 | 88.9 | 157 | 72.8 | 0.0 | 0.0 | 187.7 | 185.9 | 48.8 |
| OCDF | 362636.4 | 57713.7 | 31311 | 31745.3 | 4340.0 | 12529.5 | 54721.9 | 40057.1 | 4665.2 |
| **Total** | **4556264.5** | **737087.4** | **620844** | **672095.4** | **167525.1** | **253526.8** | **1425607.2** | **1157066.3** | **187897.9** |

**Table 5C** Concentration (pg WHO-TEQ g^-1^) of individual congeners in the lower 25cm of the sediment cores

| **Congener** | **Source** | **Site 2** | **Site 3** | **Site 4** | **Site 5** | **Site 6** | **Site 7** | **Site 8** | **Site 9** |
| --- | --- | --- | --- | --- | --- | --- | --- | --- | --- |
| 2,3,7,8-TCDD | 330.4 | 142.5 | 78.9 | 95.5 | 27.1 | 110.8 | 94.9 | 102.9 | 30.5 |
| 1,2,3,7,8-PCDD | 68.2 | 45.0 | 0.0 | 44.2 | 41.8 | 62.2 | 0.0 | 0.0 | 0.0 |
| 1,2,3,4,7,8-HexCDD | 58.3 | 30.1 | 0.0 | 0.0 | 0.0 | 0.0 | 0.0 | 0.0 | 0.0 |
| 1,2,3,6,7,8-HexCDD | 0.0 | 0.0 | 0.0 | 46.7 | 30.9 | 58.8 | 40.1 | 42.0 | 28.6 |
| 1,2,3,7,8,9-HexCDD | 20.0 | 0.0 | 9.3 | 10.2 | 5.9 | 14.2 | 0.0 | 7.8 | 6.1 |
| 1,2,3,4,6,7,8-HepCDD | 596.2 | 110.3 | 212.2 | 95.5 | 23.8 | 135.3 | 59.6 | 51.1 | 20.8 |
| OCDD | 528.5 | 200.3 | 169.3 | 188.6 | 71.3 | 330.2 | 122.1 | 108.1 | 53.6 |
| 2,3,7,8-TCDF | 5.8 | 3.9 | 0.0 | 4.3 | 0.0 | 0.0 | 0.0 | 0.0 | 0.0 |
| 1,2,3,7,8-PCDD | 15.7 | 0.0 | 0.0 | 0.0 | 0.0 | 0.0 | 0.0 | 0.0 | 0.0 |
| 2,3,4,7,8-PCDF | 19.9 | 10.2 | 12.3 | 11.6 | 0.0 | 94.7 | 0.0 | 0.0 | 0.0 |
| 1,2,3,4,7,8-HexCDF | 62.7 | 2.0 | 15.2 | 16.1 | 7.3 | 25.9 | 11.5 | 19.2 | 9.2 |
| 1,2,3,6,7,8-HexCDF | 55.4 | 2.0 | 2.0 | 2.0 | 2.0 | 2.0 | 2.0 | 2.0 | 2.0 |
| 2,3,4,6,7,8-HexCDF | 216.0 | 2.0 | 2.0 | 2.0 | 2.0 | 2.0 | 2.0 | 2.0 | 2.0 |
| 1,2,3,7,8,9-HexCDF | 2.6 | 2.6 | 3.2 | 2.0 | 2.0 | 2.0 | 2.0 | 2.0 | 2.0 |
| 1,2,3,4,6,7,8-HepCDF | 2.0 | 2.0 | 2.0 | 9.9 | 5.5 | 12.2 | 7.0 | 6.5 | 4.0 |
| 1,2,3,4,7,8,9-HepCDF | 3.4 | 2.0 | 1.8 | 0.7 | 0.0 | 1.9 | 1.3 | 0.8 | 0.0 |
| OCDF | 43.2 | 17.3 | 9.2 | 9.5 | 3.8 | 12.0 | 5.5 | 5.1 | 1.1 |
| **Total** | 2454.8 | 616.3 | 565.1 | 597.2 | 246.0 | 1016.8 | 379.5 | 387.0 | 180.1 |

**Table 5D** Concentration (pg WHO-TEQ g^-1^) of individual congeners in the lower 25cm of the sediment cores

| **Congener** | **Source** | **Site 2** | **Site 3** | **Site 4** | **Site 5** | **Site 6** | **Site 7** | **Site 8** | **Site 9** |
| --- | --- | --- | --- | --- | --- | --- | --- | --- | --- |
| 2,3,7,8-TCDD | 661.1 | 149.4 | 86.7 | 0.0 | 159.6 | 24.9 | 117.0 | 45.0 | 0.0 |
| 1,2,3,7,8-PCDD | 55.8 | 49.1 | 41.4 | 31.8 | 0.0 | 38.5 | 0.0 | 0.0 | 0.0 |
| 1,2,3,4,7,8-HexCDD | 172.2 | 0.0 | 0.0 | 2.3 | 0.0 | 2.5 | 0.0 | 0.0 | 0.0 |
| 1,2,3,6,7,8-HexCDD | 173.6 | 0.0 | 0.0 | 28.6 | 26.9 | 11.9 | 52.1 | 52.8 | 31.5 |
| 1,2,3,7,8,9-HexCDD | 30.4 | 15.0 | 11.3 | 4.5 | 13.0 | 4.5 | 9.2 | 7.6 | 0.0 |
| 1,2,3,4,6,7,8-HepCDD | 580.6 | 112.1 | 102.9 | 16.4 | 178.5 | 10.1 | 96.7 | 110.6 | 14.1 |
| OCDD | 1232.4 | 255.2 | 173.6 | 48.2 | 405.2 | 12.4 | 212.7 | 222.4 | 54.3 |
| 2,3,7,8-TCDF | 9.4 | 4.6 | 3.6 | 0.0 | 0.0 | 3.0 | 0.0 | 0.0 | 0.0 |
| 1,2,3,7,8-PCDD | 7.8 | 8.8 | 0.0 | 0.0 | 0.0 | 1.5 | 0.0 | 0.0 | 2.0 |
| 2,3,4,7,8-PCDF | 685.7 | 0.0 | 0.0 | 0.0 | 0.0 | 7.3 | 0.0 | 0.0 | 0.0 |
| 1,2,3,4,7,8-HexCDF | 102.7 | 24.9 | 12.8 | 2.0 | 23.8 | 4.3 | 25.0 | 17.4 | 2.0 |
| 1,2,3,6,7,8-HexCDF | 638.4 | 2.0 | 2.0 | 2.0 | 2.0 | 3.0 | 2.0 | 2.0 | 2.0 |
| 2,3,4,6,7,8-HexCDF | 680.3 | 2.0 | 2.0 | 2.0 | 2.0 | 2.3 | 2.0 | 2.0 | 2.0 |
| 1,2,3,7,8,9-HexCDF | 7.3 | 2.0 | 5.3 | 2.0 | 2.0 | 2.3 | 2.0 | 2.0 | 2.0 |
| 1,2,3,4,6,7,8-HepCDF | 54.7 | 2.0 | 2.0 | 3.5 | 15.4 | 1.4 | 8.0 | 9.6 | 3.9 |
| 1,2,3,4,7,8,9-HepCDF | 5.4 | 0.4 | 1.6 | 0.0 | 1.9 | 0.3 | 0.0 | 1.6 | 0.5 |
| OCDF | 108.8 | 18.0 | 9.4 | 1.3 | 16.4 | 0.2 | 8.0 | 9.4 | 1.4 |
| **Total** | 7507.1 | 710.2 | 493.1 | 157.5 | 910.0 | 155.8 | 581.7 | 526.5 | 131.5 |

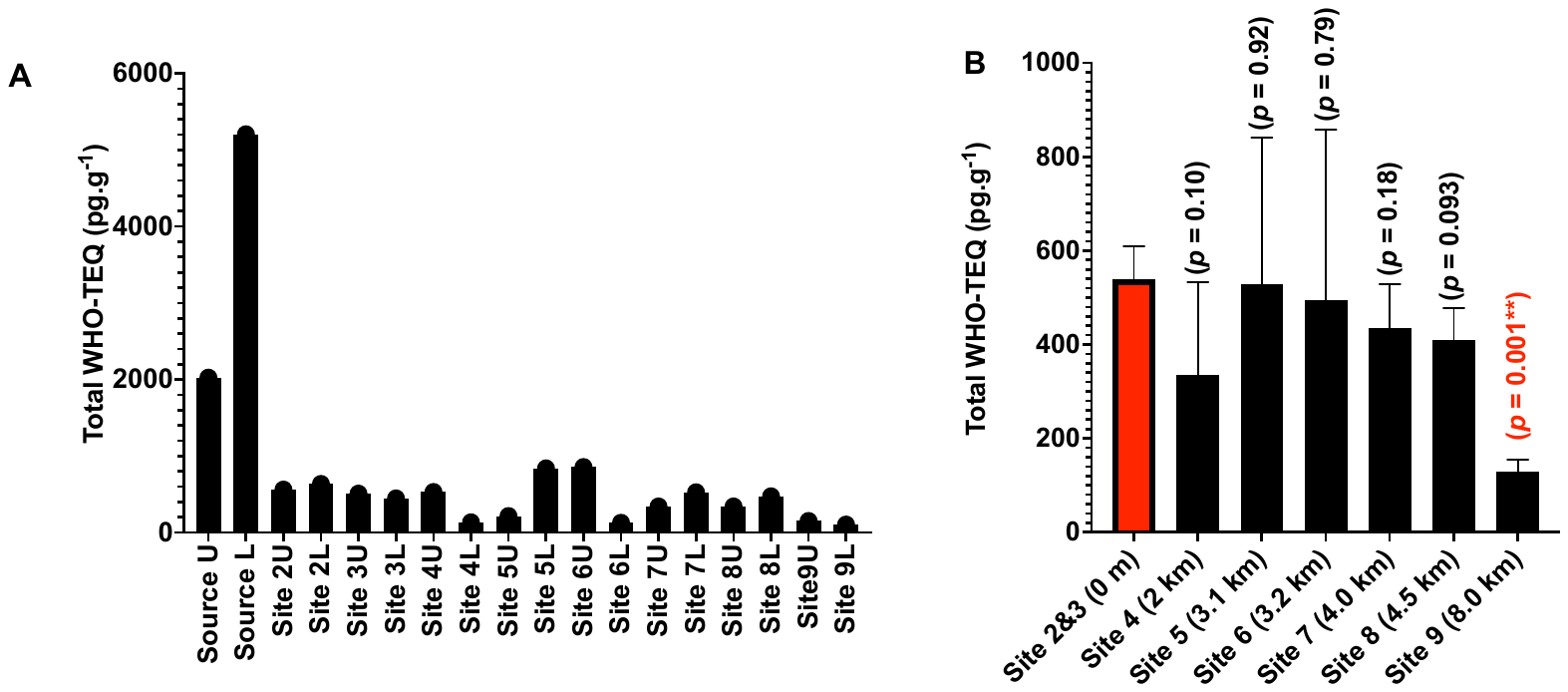

**Figure 2: Panel A**: PCDD/F concentrations at all sampling locations at both depths (Upper (U) and Lower (L). **Panel B:** the mean of PCDD/F upper and lower PCDD/F concentrations at all locations (excluding the source). *P*-values are derived from *t*-test comparison of concentrations at each location with Homebush Bay sites (i.e. sites 2 & 3). Error bars represent standard deviation from the mean (n = 3)

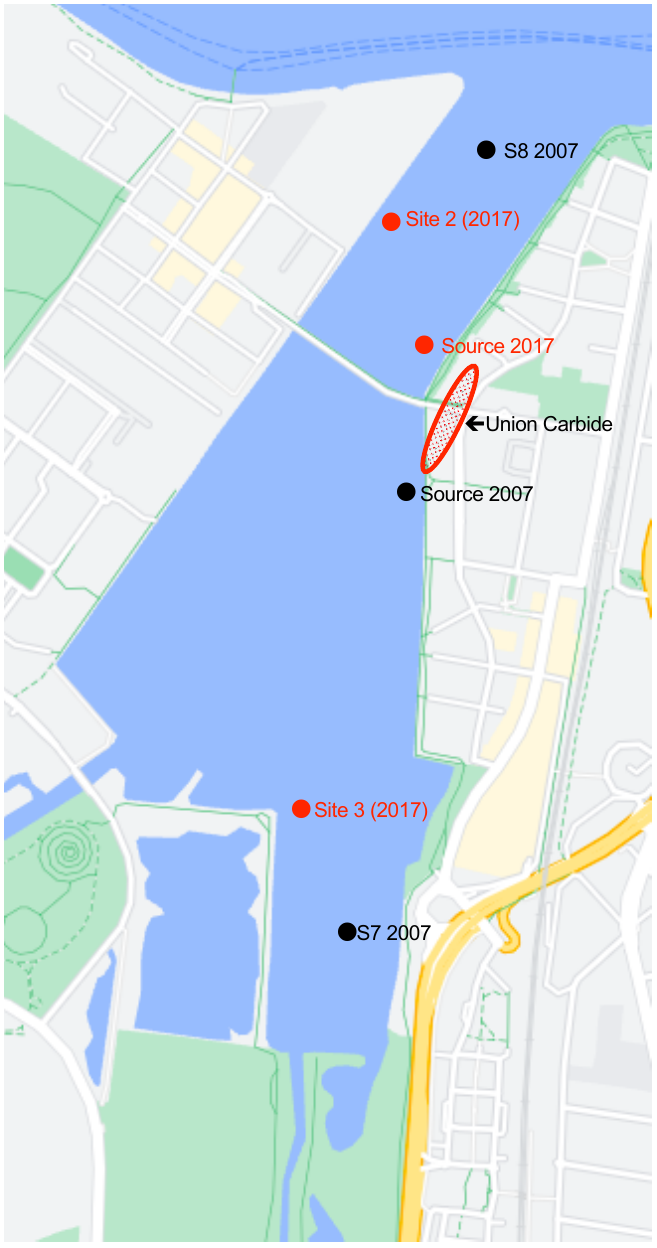

**Figure S3.** Precise sample locations of sediments sampled from Homebush Bay in 2007 and 2017, and the location of the Union Carbide facility on the Eastern shore of Homebush Bay (Rhodes Peninsula).

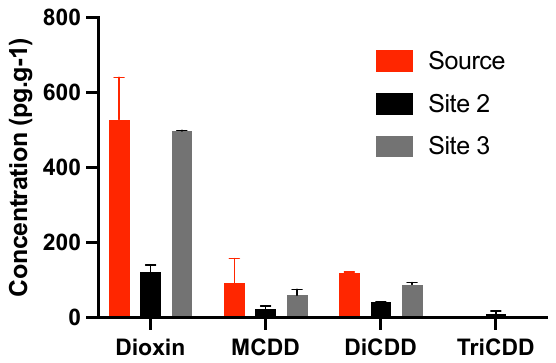

**Figure S4.** 2,3,7,8-TCDD dechlorination products in Homebush Bay sediments. Error bars represent the standard deviation from the mean (n = 2).
